# Interleukin-13-mediated alterations in esophageal epithelial mitochondria contribute to tissue remodeling in eosinophilic esophagitis

**DOI:** 10.1101/2025.04.02.646853

**Authors:** Jazmyne L. Jackson, Reshu Saxena, Mary Grace Murray, Abigail J. Staub, Alena Klochkova, Travis H. Bordner, Courtney Worrell, Annie D. Fuller, John M. Crespo, Andres J. Klein-Szanto, John Elrod, Tatiana A. Karakasheva, Melanie Ruffner, Amanda B. Muir, Kelly A. Whelan

## Abstract

**Background:** The significance of mitochondria in EoE pathobiology remains elusive.

**Objective:** To determine the impact of EoE inflammatory mediators upon mitochondrial biology in esophageal epithelium, the mechanisms mediating these effects, and their functional significance to EoE pathobiology.

**Methods:** Mitochondria were evaluated in human biopsies, MC903/Ovalbumin-induced murine EoE, and human esophageal keratinocytes. Esophageal keratinocytes were treated with EoE-relevant cytokines and JAK/STAT inhibitor ruxolitinib. To deplete mitochondria, 3D organoids generated from *TFAM^loxp/loxp^* mice were subjected *ex vivo* to Cre or siRNA against Transcription factor A, mitochondria (TFAM) was transfected into esophageal keratinocytes. Mitochondrial respiration, membrane potential, and superoxide levels were measured.

**Results:** We find evidence of increased mitochondria in esophageal epithelium of patients with EoE and mice with EoE-like inflammation. In esophageal keratinocytes, IL-4 and IL-13 increase mitochondrial mass. IL-13 increases mitochondrial biogenesis in a JAK/STAT-dependent manner. In 3D organoids, IL-13 limits squamous cell differentiation (SCD), and this is blunted upon TFAM depletion. IL-13 decreases mitochondrial respiration and superoxide level, although mitochondria remain intact. IL-13-mediated suppression of superoxide was abrogated upon TFAM depletion in esophageal keratinocytes.

**Conclusions:** We report that increased mitochondrial mass is a feature of EoE. Among EoE-relevant cytokines, IL-13 is the primary driver of increased mitochondrial mass in esophageal keratinocytes by promoting mitochondrial biogenesis in a JAK/STAT-dependent manner. IL-13-mediated accumulation of mitochondria impairs SCD in esophageal keratinocytes and also suppresses oxidative stress, a factor that is known to induce SCD. These findings identify a novel mechanism through which IL-13 promotes EoE-associated epithelial remodeling.

**Clinical Implication:** These findings further lay a foundation for exploration of level of esophageal epithelial mitochondria as a predictive biomarker for response to dupilumab.

**Capsule summary:** IL-13 promotes mitochondrial biogenesis in esophageal epithelium, contributing to impaired squamous cell differentiation.

## Introduction

Eosinophilic esophagitis (EoE) is a chronic food allergen- and immune-mediated disease that exerts a significant clinical and financial burden worldwide^1^. EoE is characterized by esophageal eosinophilia and tissue remodeling in esophageal epithelial, muscle, and stromal compartments^1, 2^. EoE symptoms include vomiting, dysphagia and food impaction, all of which negatively impact patient quality of life^3–7^. While dietary elimination and corticosteroids are therapeutic mainstays in EoE, the monoclonal antibody dupilumab, which targets interleukin (IL)-4 receptor (IL-4R)α, became the first and only FDA-approved targeted therapy for use in EoE patients in 2022^8, 9^. This significant advance in EoE patient care was guided by experimental evidence supporting the importance of T helper (Th)2 signaling in EoE. Presently, our understanding of the molecular mechanisms through which Th2 cytokines drive EoE pathobiology remain incompletely understood.

In response to allergens, esophageal epithelial cells secrete cytokines, including thymic stromal lymphopoietin^10, 11^, to recruit lymphocytes that then produce Th2 cytokines IL-4, IL-5, and IL-13. In esophageal epithelium, IL-4 and IL-13 bind to IL-4R to promote JAK-mediated phosphorylation of signal transducer and activator of transcription (STAT) proteins^12^. Phosphorylated STATs then dimerize and translocate to the nucleus where they bind DNA and direct target gene transcription^13^. IL-4R-mediated signaling in esophageal epithelium is a key driver of EoE with targeted genetic depletion of epithelial IL-4Rα subunit protecting mice from experimental EoE *in vivo*^14^, supporting IL-4R-mediated signaling in esophageal epithelium as a key driver of EoE. Although epithelial-associated IL-4 and IL-13-responsive genes, including eotaxin-3^15, 16^, have been linked to EoE pathobiology, studies in mice suggest a predominate role for IL-13 in experimental EoE^14^. We have demonstrated that IL-13 promotes epithelial remodeling, including basal cell accumulation and impaired differentiation, in 3D esophageal organoids^17–19^ and these findings have been recapitulated in murine models^14, 20–22^. Additionally, IL-13-mediated induction of calpain-14^23^, synaptopodin^24^ and follistatin^21^ contribute to impaired epithelial squamous cell differentiation (SCD) *in vitro*.

Oxidative stress contributes to regulation of esophageal epithelial differentiation with excessive reactive oxygen species (ROS) promoting SCD^25^. Moreover, mechanisms that suppress oxidative stress have been linked to EoE-associated epithelial remodeling^18, 21, 26^. For example, our own work has demonstrated that autophagic flux is induced in esophageal keratinocytes responding to IL-13 or tumor necrosis factor (TNF)α to limit oxidative stress^18^. Mitochondria serve as a critical source of cellular ROS, which is produced as a byproduct of oxidative phosphorylation (OXPHOS). Genetic evidence has identified a potential role for mitochondria in the pathogenesis of EoE^27^. Whole exome sequencing on patients with EoE and unaffected family members revealed enrichment in rare, damaging variants in two genes: *DHTKD1 (*Dehydrogenase and Transketolase Domain Containing 1), which contributes to mitochondrial lysine metabolism and ATP production^28, 29^, and its homolog *OGDHL (*Oxoglutarate Dehydrogenase-Like)^27^. *In vitro* experiments further revealed significant reductions in mitochondrial respiration in esophageal fibroblasts derived from patients with EoE expressing *DHTKD1* or *OGDHL* variants^27^. Impaired mitochondrial respiration was also identified upon DHTKD1 knockdown in normal esophageal epithelial cultures that further displayed increased levels of ROS^27^. Recently, a study identified evidenced of metabolic dysfunction, including enrichment of OXPHOS pathway, using transcriptomic data from mucosal biopsies of patients with EoE^30^.

Here, evaluate the impact of EoE-associated inflammatory mediators upon mitochondrial biology in esophageal epithelium and exploring the mechanisms contributing to these effects and their functional significance in EoE pathobiology.

## Methods

### Human Subjects

Human subjects were subjected to endoscopic biopsy collection by esophagogastroduodenoscopy (EGD) during routine clinical care under protocols approved by the Institutional Review Board at Hospital of the University of Pennsylvania. At the time of diagnostic esophagogastroduodenoscopy, pinch biopsies were obtained for clinical evaluation. Written informed consent was obtained from each subject or their legal guardian (where appropriate). Patients who met clinical criteria for EoE^31^, with mucosal eosinophils per high power field (eos/HPF) were classified as active EoE. Inactive EoE patients had a history of active EoE and clinical showed resolution of esophageal eosinophilia (<15 eos/HPF) at the time of follow up EGD. Normal subjects included those reporting symptoms warranting EGD, but who demonstrated no endoscopic or histological abnormalities in the upper gastrointestinal tract. Subjects with a history of inflammatory bowel disease, celiac disease, GI bleeding or any other acute or chronic intestinal disorders were excluded from recruitment. Demographic and clinical information on human subjects whose esophageal biopsies were evaluated in the current study is provided in **Table E1 in the Online Repository**.

### Murine Studies

All animals were purchased, bred, and treated under Temple University Institutional Animal Care and Use Committee-approved protocols. EoE was induced in wild type C57/B6 mice using the MC903/Ovalbumin (OVA) protocol^18, 32^ with the following modifications: sensitization period during which MC903 (2 nmol; Tocris Bioscience, Cat# 2700) and OVA (100 μg; Sigma-Aldrich, Cat# A5503-50G) are applied to ears was shortened to from 14 days to 12 days; no treatments were administered on days 13 and 14; OVA in drinking water was increased from 1.5 g/mL to 15 g/mL; challenge period was extended from 4 days to 18 days. Mice treated with MC903 only were used as controls. **Figure E1A in the Online Repository** provides a schematic of the protocol used. B6.Cg-Tfam^tm1.1Ncdl/^J mice (Cat# 026123) were purchased from Jackson Laboratories at age 4-8 weeks. All purchased mice were allowed to acclimate for at least 2 weeks prior to use in experiments. Following experimental protocols, esophagi were harvested and processed as described below.

### Immunohistochemistry (IHC) & Histological Evaluation

Formalin-fixed paraffin-embedded tissue section on glass slides were stained with the hematoxylin & eosin (H&E) or the following primary antibodies as previously described^18^: MTCO1 [1D6E1A8] (1:250; Abcam, Cat# ab14705) and TFAM (1:500; Sigma Cat# SAB1401382-50UG). A pathologist blinded to clinical parameters (AKS) scored MTCO1 staining on scale from 0 (negative) to 3, taking into account stain intensity and distribution.

### Cell Culture

The immortalized normal esophageal keratinocyte cell line EPC2-hTERT was provided as a generous gift from Drs. Anil Rustgi and Hiroshi Nakagawa (Columbia University). Primary cell cultures generated from esophageal biopsies from a normal human subject (EPC203) and an active EoE patient (EPC128) were provided as a generous gift from Dr. Amanda Muir (CHOP). All cells were cultured in keratinocyte serum free media (KSFM; Gibco, Cat# 17005042) supplemented with recombinant epidermal growth factor (1ng/mL), bovine pituitary extract, (50 μg/mL), and penicillin/streptomycin (1% v/v, Gibco, Cat# 15140-122), as previously described^18^. For experiments with TNFα (R&D Systems, Cat# 210-TA-005/CF; 40 ng/mL), IL-4 (R&D Systems, Cat# 204-IL-010; 10 ng/mL), IL-5 (R&D Systems, Cat# 205-IL-005; 10 ng/mL), IL-13 (R&D Systems, Cat# 213-ILB-005/CF; 10 ng/mL) or IL-1β (R&D Systems, Cat# 201-lB-005; 10 ng/mL), 30,000 cells were plated in 6-well plates, or 100,000 cells were plated on 10-cm plates 24 hours prior to stimulation with cytokines, then media was changed every 48 hours for a total of 7 days. 10 ng/μl stocks of cytokines (20 ng/μl of TNFα) were dissolved in 0.1% Bovine Serum Albumin (Fisher, Cat# BP9703-100) in 1X D-PBS (Gibco, Cat# 14190-144).

To deplete TFAM in esophageal keratinocytes, small interfering RNA oligonucleotides targeting TFAM (Dharmacon, ID: J-019734-06) or a non-targeting (NT) siRNA pool (ID: D-001810-10-05) were diluted in 500 μL Opti-MEM reduced serum media (Gibco, Cat# 31985088) to a final concentration of 10 nM per well of a 6 well tissue culture dish then 5 μL of Lipofectamine™ RNAiMAX transfection reagent (ThermoFisher Scientific, Cat# 13778150) was added to each well. Plates were incubated at room temperature for 20 minutes then 300,000 EPC2-hTERT cells diluted in 2 mL KSFM without antibiotics were added to each well. 72 hours after siRNA transfection, cells were utilized for experiments.

To suppress the JAK-STAT6 pathway, the JAK inhibitor, Ruxolitinib (100 ng/mL; Invivogen, Cat# tlrl-rux-3) was utilized. The inhibitor was refreshed every 48-72 hours throughout the 7-day period of cytokine stimulation.

### Immunoblotting

Whole cell lysates from esophageal keratinocyte cultures or tissue lysates from peeled murine esophageal epithelium were subjected to immunoblotting as described previously^18, 33^ using the following primary antibodies: MTCO1 (Abcam, Cat# ab14705, 1:1000), TFAM (Cell Signaling, Cat# 7495, 1:1000), phospho-Stat6 Tyrosine 641 (Cell Signaling, Cat#), STAT6 (Cell Signaling, Cat#), and Actin (Invitrogen, Cat# MA1-744, 1:10000) . Targeted proteins were visualized using chemiluminescence detection reagents (ProSignal Femto ECL Reagent, Cat # 20-302) and imaged on an iBright imaging system (Invitrogen).

### Quantitative Polymerase Chain Reaction (qPCR)

For mitochondrial DNA (mtDNA) assays^34^, DNA isolation was performed on esophageal keratinocyte cells or peeled murine esophageal epithelium using DNeasy Blood and Tissue Kit (Qiagen, Cat# 69506) according to the manufacturer’s instructions. DNA concentration was determined using Qubit™ dsDNA HS Assay Kit according to the manufacturer’s instructions (Invitrogen, Cat# Q32851). qPCR was performed using PowerUp™ SYBR™ green master mix (ThermoFisher Scientific, Cat# A25743).

To assess mtDNA, primers for mtDNA D-Loop (murine), *MTCO1* (human) and *ND6* (human) were used. To assess nuclear DNA, primers for *Ikb*β (murine), *COXIV* (human), and *GAPDH* (human) were used. The relative fold change between the noted mtDNA-encoded genes and nuclear DNA-encoded genes (*Ikb*β or *GAPDH*) was calculated using the delta delta CT method. In human specimens, the relative expression of the nuclear DNA-encoded gene *COXIV* was used as an additional internal control.

For reverse transcriptase PCR, RNA extraction was performed with RNeasy Mini Kit (Qiagen, Cat# 74106) according to manufacturer’s instructions. RNA concentration was measured using Qubit™ RNA HS Assay Kit (Invitrogen, Cat# Q32852). Reverse transcription was performed using the High-Capacity cDNA Reverse Transcription Kit for RT-qPCR (Thermo Fisher Scientific, Cat# 4368814). qPCR was performed using PowerUp™ SYBR™ green master mix (Thermo Fisher Scientific, Cat# A25743). Primers for *TFAM, MFN1, MFN2, DRP1, PINK1, PARKIN,* and β*-Actin* were used. The relative levels of *TFAM, MFN1, MFN2, DRP1, PINK1,* or *PARKIN* relative to β*-*Actin was calculated using the delta delta CT method. All primer sequences are listed in **Table E2 in the Online Repository**.

### Live Cell Imaging

Mitochondria were measured using MitoTracker Green dye (Invitrogen, Cat# M7514). Superoxide level was measured using MitoSOX Red (Invitrogen, Cat# M36008). Esophageal keratinocytes were grown in 100 mm tissue culture plates in the presence or absence of IL-13 (10 ng/mL) for 7 days. On the 6^th^ day, cells were plated on 30 mm glass bottom collagen coated plates overnight then stained with indicated dyes according to the manufacturer’s protocol. In brief, cells were stained with KSFM containing 200 nM MitoTracker Green or 5 μM of MitoSOX Red and counterstained with 2 µM of Hoechst (Invitrogen, Cat# 33342), for 30 min at 37°C, 5% CO_2_. After staining, cells were washed 3X with HBSS and live cells were analyzed by Leica Confocal Microscope using a 63X oil objective. Cells were maintained at 37°C, 5% CO_2_ during imaging. Images were quantified using ImageJ software. Mean integrated intensities of images (from twenty randomly chosen fields) after background subtraction were interpreted as a quantitative measure of mitochondria.

### Oxygen Consumption Rate (OCR) Assay

Cellular oxidative phosphorylation (OXPHOS) was monitored using a Seahorse Bioscience Extracellular Flux Analyzer (XF96e, Seahorse Bioscience) by real time OCR measurement following the protocol of XF Cell Mito Stress Kit (Agilent Technologies, Cat# 103015-100). In brief, EPC2-hTERT cells were grown in 100 mm tissue culture plates with 10 ng/mLIL-13 treatment for 7 days. On the 6^th^ day, 100,000 cells/well were seeded onto XF96 96-well plates in 200 μL of growth medium and incubated overnight at 37 °C, 5% CO_2._ One hour prior to assay, culture media was replaced by XF assay media (180 μL; Agilent Technologies, Cat# 102353-100) and cells were incubated at 37°C incubator without CO_2_ regulation for 1 hour to allow to pre-equilibration with the XF assay medium which was further supplemented with 25 mM glucose (Sigma, Cat# D8270) and 1 mM sodium pyruvate (pH 7.4; Sigma, Cat# S8636). Cells were sequentially exposed to oligomycin (1 μM), carbonilcyanide p-triflouromethoxyphenylhydrazone (FCCP; 3 μM) and Rotenone plus Antimycin A (1 μM). Basal OCR levels were recorded followed by OCR levels after the injection of individual compounds that inhibit respiratory mitochondrial electron transport chain (ETC) complexes. Results were normalized according to cell protein concentration. Sequential addition of oligomycin, FCCP, as well as rotenone & antimycin A allowed an estimation of the contribution of individual parameters for basal respiration, proton leak, maximal respiration, spare respiratory capacity, non-mitochondrial respiration, and ATP production.

### ATP Assay

ATP production was assessed by using the CellTiter-Glo 2.0 Kit (Promega, Cat# G9241) according to the manufacturer’s protocol. In brief, EPC2-hTERT cells were grown in 100 mm tissue culture plates in the presence or absence of IL-13 (10 ng/mL) for 7 days. On the 6^th^ day, 100,000 cells/well were seeded onto 96-well solid white plates in 100 μL of growth medium per well and incubated overnight with at 37 °C, 5% CO_2_ . 100µL of CellTiter-Glo® 2.0 Reagent was added to 100 µl of medium containing cells, followed by mixing for 2 minutes and incubation at room temperature for 10 minutes to stabilize the luminescent signal. Luminescence was measured by using Promega GloMax-Multi+ detection system. Results were expressed as percentage decrease from untreated sample.

### Flow cytometry

EPC2-hTERT cells were treated with IL-13 (10 ng/mL) for 7 days with media changes occurring every 48 hours. Mitochondrial membrane potential (depolarized mitochondria) was determined by staining cells with MitoTracker Green (ThermoFisher Scientific, Cat# M7514) at a final concentration of 25 nM and MitoTracker Deep Red (ThermoFisher Scientific, Cat# M22426) at a final concentration of 100 nM. Mitochondrial superoxide production was measured by staining cells with MitoSOX Red (ThermoFisher Scientific, Cat# M36008) at a final concentration of 5 µM. Cells were stained with each dye for 30 minutes at 37°C. Cells were washed with and resuspended in 1X HBSS (Gibco, Cat# 14175095). Flow cytometry was performed using Symphony A5 flow cytometer (BD Biosciences) and data were analyzed with FlowJo version 10.0.

### 3D Esophageal Organoid Assays

Whole esophagi were dissected from 2 female, 2 male B6.Cg-Tfam tm1.1Ncdl/J mice (age range 12-24 weeks old). Esophageal epithelium was physically separated from underlying submucosa using forceps then the esophagus was cut open longitudinally to expose the epithelial surface. Peeled epithelium was incubated in 1 mL 1X Dispase (Corning, Cat# 354235) in HBSS (Gibco, Cat# 14025-076) for 10 minutes at 37°C with shaking at 1000 RPM. Following removal from Dispase, esophageal epithelium was chopped into 3 pieces with sharp scissors then incubated in 1 mL of 0.25% Trypsin-EDTA (Genesee Scientific, Cat# 25-510) for 10 minutes at 37° C with shaking at 1000 RPM. Trypsin and tissue pieces were forced through a cell strainer (70 μm) into a 50 mL conical tube containing 4 mL of 250µg/mL soybean trypsin inhibitor (Gibco, Cat# 17975-029) in 1X PBS. Cells were pelleted at 1000 RPM for 5 minutes then resuspended in 500 μL of complete ‘mouse’ KSFM without calcium chloride (Gibco, Cat# 10725-018) supplemented with recombinant epidermal growth factor (1 ng/mL), bovine pituitary extract, (50 mg/mL), and penicillin/streptomycin (1% v/v; Gibco, Cat# 15140-122), and CaCl2 to a final concentration of 0.018 mM (Acros Organics, Cat# 349610025) as previously described^35^. Cell number and viability were measured by Automated Cell Count (Invitrogen, Countess II FL). Single cell suspension in mouse KSFM medium was mixed with 90% Matrigel (Corning, Cat# 354234) to initiate 3D organoid formation. Using 24 well plates, 4,000 cells were seeded per well in 50 μL Matrigel. After solidification, 500 μL of Advanced DMEM/F12 (Gibco, Cat# 12634-101) supplemented with 1X Glutamax (Gibco, Cat# 35050-061), 1mM HEPES (Gibco, Cat# 15630-080), 1% v/v penicillin-streptomycin (Gibco, Cat# 15140-122), 1X N2 Supplement (Gibco, Cat# 17502-001), 1X B27 Supplement (Gibco, Cat# 17504-044), 0.1 mM N-acetyl-L-cysteine (Fisher Chemical, Cat# 616-91-1), 50 ng/mL human recombinant EGF (Gibco, Cat# 10450-013), 2.0% Noggin/R-Spondin-conditioned media and 10 μM Y27632 (Tocris, Cat# 1254) were added to each well. To initiate recombination, Ad5CMVeGFP virus (1*10^11^pfu/mL; Cat# ad4327) or Ad5CMVCre-eGFP virus (1*10^11^ pfu/mL; Cat# ad4216) purchased from University of Iowa Viral Vector Core was added to growth media at a 1:100 dilution. At the time of plating, organoids were also treated with IL-13 (10 ng/mL) or vehicle. After 48 hours, media was changed to remove virus. Treatment with IL-13 (10 ng/mL) was continued until day 7 with media changes occurring every other day. Organoids were grown for 7 days before recovering from Matrigel with Dispase I and fixing overnight in 4.0% paraformaldehyde. Specimens were embedded in 2.0% Bacto-Agar: 2.5% gelatin prior to paraffin embedding.

### EGID Database

To assess the expression of mediators of mitochondrial biology in esophageal biopsies, we accessed the “EoE Transcriptome by RNA Sequencing^36^” section on the EGIDExpress database (https://egidexpress.research.cchmc.org/data/), a site that contains data relating to Eosinophilic Gastrointestinal Diseases (EGIDs).

### Statistics

Descriptive statistics are presented as mean ± standard error of the mean (SEM). Student’s t-test or one-way analysis of variance was used to determine significance for comparison of two groups. For groups of three or larger, One-way ANOVAs followed by Dunnett’s or Tukey’s multiple comparisons were performed using GraphPad Prism version 9.0.2 for macOS (GraphPad Software, San Diego, California USA). p<0.05 was considered statistically significant.

## Results

### Evidence of increased mitochondria in esophageal epithelium of EoE patients and mice with EoE-like inflammation

To explore the impact of EoE inflammation upon mitochondria, we evaluated expression of mitochondrial proteins by IHC in human patients with EoE and control subjects with normal esophageal pathology. Staining for MTCO1 (mitochondrially encoded cytochrome c oxidase I), a mitochondrially-encoded subunit of Complex IV of the ETC, was significantly increased in patients with active EoE inflammation as compared to either normal controls or patients with inactive EoE (**Fig. 1A, B**). Increased MTCO1 expression at the protein level was also found in mice with EoE-like inflammation induced by MC903/OVA (**Fig. 2A**), a food allergen-mediated model that exhibits features consistent with EoE pathophysiology as found in humans, including eosinophil-rich inflammatory infiltrates in esophageal mucosa (**see Fig E1A, B in the Online Repository**), impairment of the epithelial proliferation/differentiation gradient, and food impactions^18, 32^. As MTCO1 represents only a single mitochondrial protein, we continued to explore whether EoE inflammation impacts mitochondrial mass. Peeled esophageal epithelium of MC903/OVA-treated mice displayed a significant increase in the ratio of mtDNA relative to nuclear DNA as compared to their MC903-only treated counterparts, the latter of which develop robust atopic dermatitis but fail to exhibit esophageal inflammation (**Fig. 2B**).

**Figure 1.**
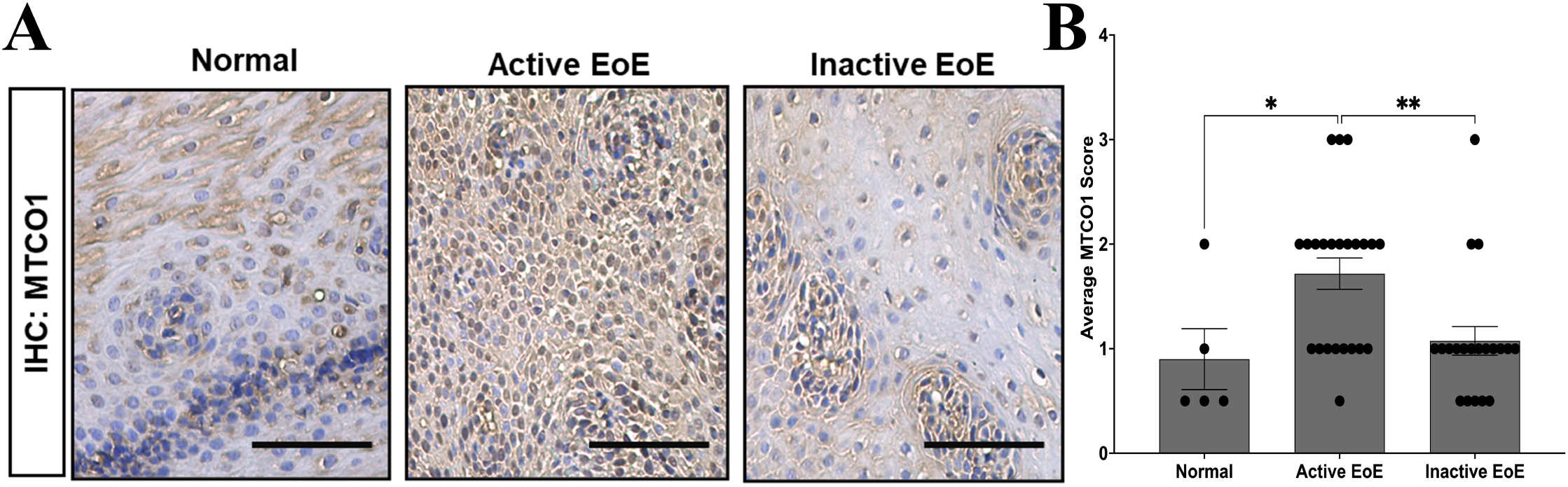
Evidence of increased mitochondria in EoE patients. (**A**) Representative images of immunohistochemistry (IHC) staining for mitochondrially-encoded cytochrome oxidase 1 (MTCO1) in esophageal biopsies from human subjects classified as normal, active EoE or inactive EoE. Scale bars, 50 µm. (**B**) Average MTCO1 score in esophageal epithelium of indicated groups. Data in bar graphs presented as mean ± SEM. *p<0.05; **p<0.01.

**Figure 2.**
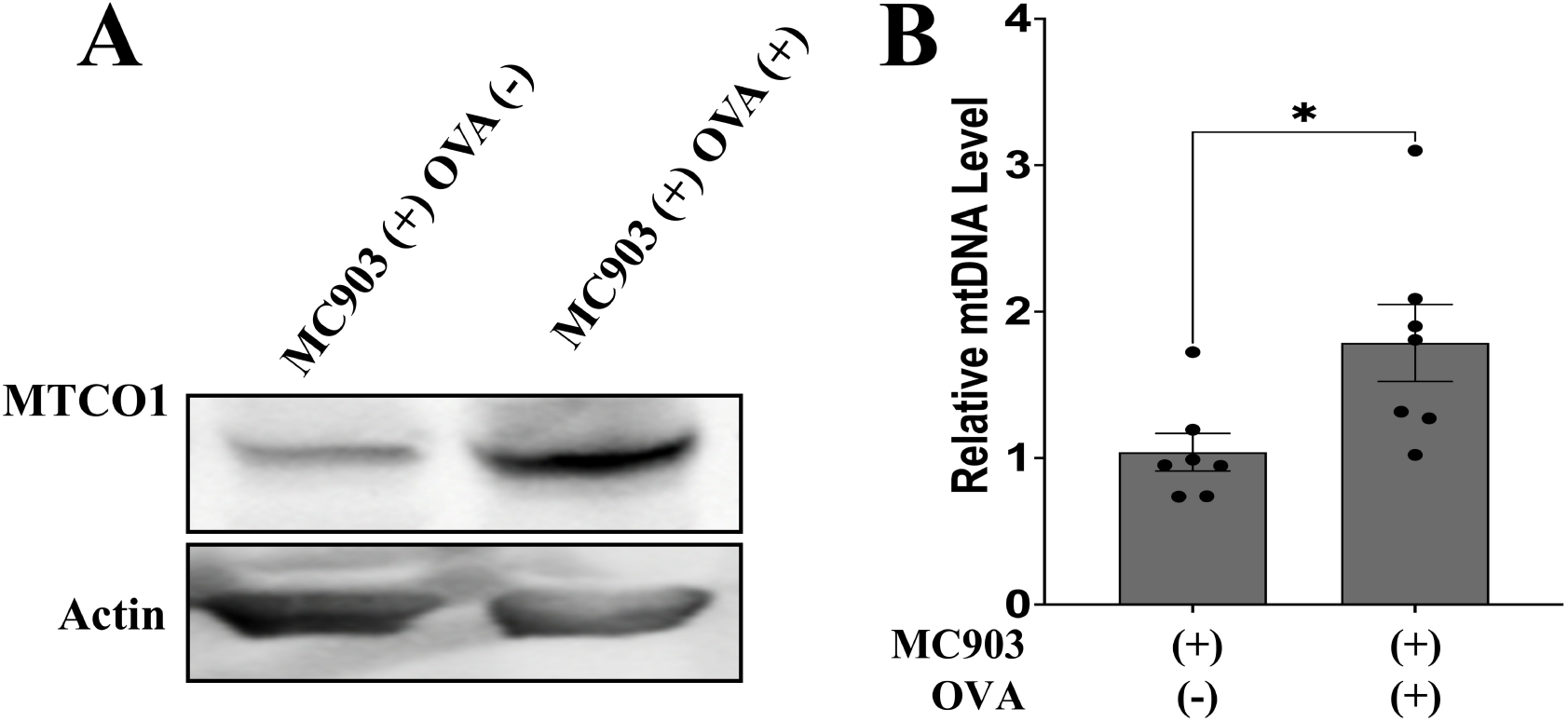
Evidence of increased mitochondria in esophageal epithelium of mice with EoE-like inflammation. C57B6 mice were treated with MC903 and Ovalbumin (OVA) over 32-days then esophagi were dissected, and epithelium was peeled. MC903-only treated mice served as controls. (**A**) Immunoblotting for MTCO1 protein expression with Actin as a loading control. (**B**) Quantitative PCR (qPCR) was used to assess mtDNA levels with *Ikb*β as an internal control. Data in bar graphs presented as mean ± SEM. *p<0.05; n=7 for MC903-only controls and n=7 for MC903+/OVA+.

### IL-13 and IL-4 increase mitochondrial mass in esophageal keratinocytes

We next aimed to define the signals in the EoE inflammatory milieu that promote increased mitochondria in esophageal epithelial cells. To that end, we stimulated the immortalized normal esophageal cell line EPC2-hTERT with a panel of EoE-relevant cytokines, consisting of IL-4, IL-5, IL-13, IL-1β, and TNFα, for 7 days. To assess mitochondria, we performed DNA qPCR for the mitochondrially encoded genes *MTCO1* and *ND6*. We also evaluated DNA level of *COXIV*, a nuclear gene that encodes the mitochondrial protein cytochrome c oxidase 4. A significant increase in both *MTCO1* and *ND6* was detected in EPC2-hTERT cells treated with IL-4 or IL-13 with the effect induced by IL-13 more pronounced (**Fig. 3A**). EoE-relevant cytokines failed to influence DNA level of *COXIV*, which is expected as EoE has not been shown to induce genetic instability. mtDNA level may result from an increase in mitochondria or an increase in the mtDNA copy number within each mitochondrion. As such, we next used the mitochondrial dye MitoTracker Green to assess mitochondria in IL-13-treated esophageal epithelial cells. Confocal imaging revealed an increase in mitochondria mass along with expansion of the tubular perinuclear mitochondrial network in EPC2-hTERT cells, primary normal, and primary EoE cells treated with IL-13 (**Fig. 3B, C; see Figure E2 in the Online Repository**).

**Figure 3.**
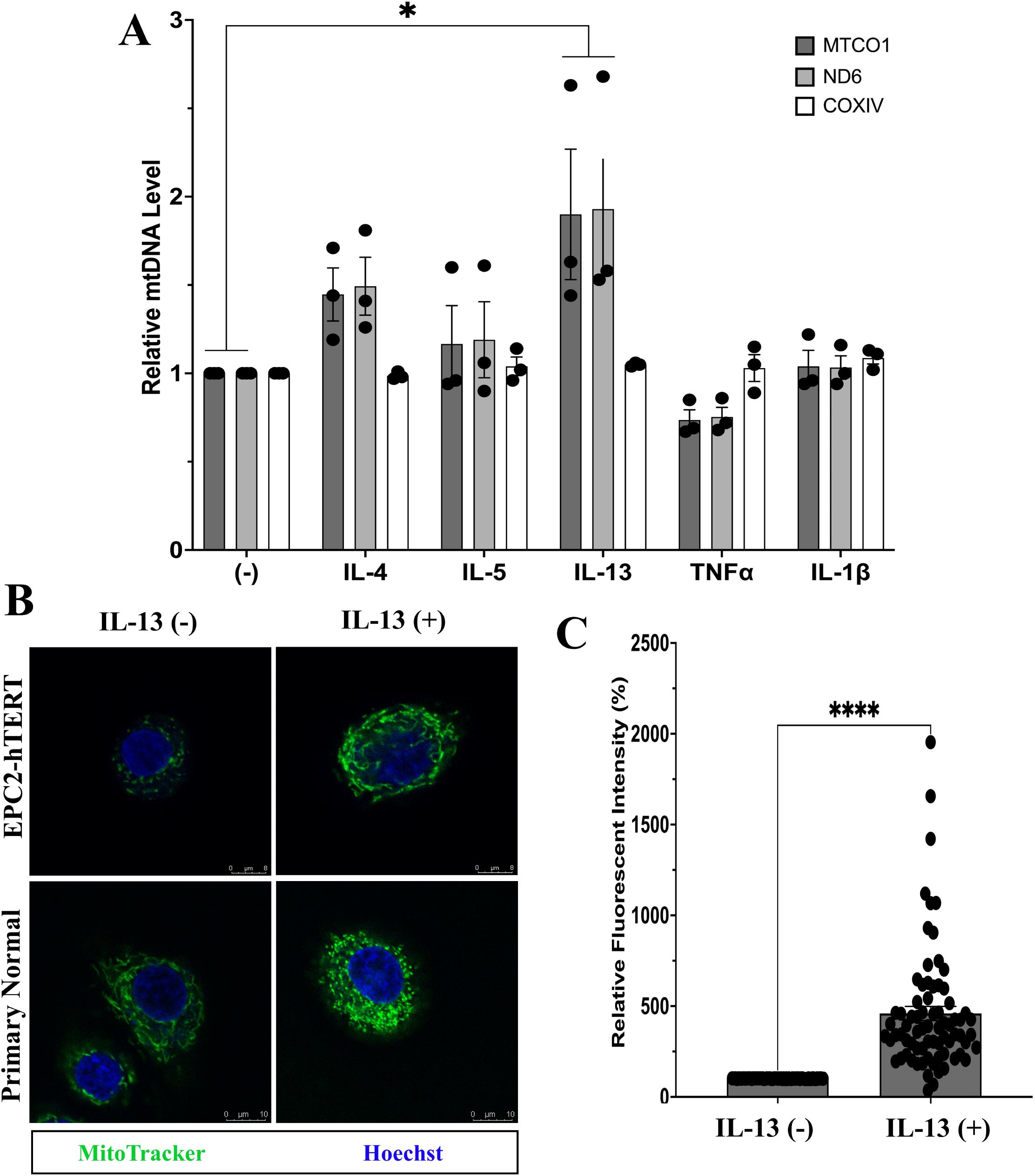
Effects of EoE-relevant cytokines on mitochondria in human esophageal keratinocytes *in vitro*. (**A**) EPC2-hTERT cells were treated with the indicated cytokines for 7 days. All interleukin (IL) treatments were at a final concentration of 10 ng/mL. Tumor necrosis factor (TNF)α treatment was at a final concentration of 40 ng/mL. qPCR evaluated levels of mitochondrially-encoded genes *MTCO1* and *ND6*, and nuclear-encoded gene *COXIV*. All genes are shown normalized to *GAPDH*. (**B, C**) EPC2-hTERT and primary esophageal epithelial cultures were treated with IL-13 (10 ng/mL) for 7 days then stained with MitoTracker Green and Hoechst 33342 to visualize mitochondria and nuclei, respectively, using confocal microscopy. (**B**) Representative images are shown. (**C**) Quantitative fluorescence intensity of MitoTracker Green-stained EPC2-hTERT cells was analyzed by ImageJ software. Data in bar graphs presented as mean ± SEM, *p<0.05; n=3.

### IL-13 and IL-4 promote expression of TFAM in esophageal keratinocytes

An increase in mitochondrial mass may be reflective of increased biogenesis, alterations in fission/fusion dynamics, or impaired mitochondrial turnover, the latter of which primarily occurs via mitophagy^37^. To determine the mechanism through which IL-13 and IL-4 treatment results in accumulation of mitochondria in esophageal keratinocytes, we next used qPCR to survey key genes involved in the regulation of each of these facets of mitochondrial biology. Both IL-13 and IL-4 induced expression of *TFAM*, a critical mediator of biogenesis^38^, in EPC2-hTERT cells. The timing of TFAM induction differed with IL-13-stimulated EPC2-hTERT cells showing significant upregulation of *TFAM* at 1, 3, and 7 days of treatment (**Fig. 4A**) while their IL-4-treated counterparts displayed induction at 1 and 5 days of treatment (**see Figure E3A in the Online Repository**). No significant expression of *MFN1* or *MFN2*, encoding critical mediators of mitochondrial fusion Mitofusion1 and Mitofusion2, or in *DRP1,* encoding the critical mediator of mitochondrial fission dynamin-related protein-1 was detected in EPC2-hTERT cells treated with either IL-13 or IL-4 (**Fig. 4B-D; see Figure E3B-D in the Online Repository**). A significant increase in expression of *PINK1* and *PARK2*, encoding mediators of mitophagy PTEN-induced kinase-1 and Parkin, was detected in EPC2-hTERT cells following 5 days of treatment with IL-13 or IL-4 (**Fig. 4E, F**; **see Figure E3E, F in the Online Repository**). We validated increased levels of TFAM protein in EPC2-hTERT cells treated with IL-13 or IL-4 for 7 days (**Fig. 4G, H**). We further assessed expression of genes involved in mitochondrial biogenesis, fission/fusion, and mitophagy in publicly available transcriptomic data from patients with active EoE and non-EoE controls (EGID database). Expression of T*FAM* was significantly increased in active EoE compared to normal controls^36^ (**Fig. 4I**). Additionally, increased levels of MFN1 and MFN2 were detected in active EoE patients which further exhibited diminished levels of *PINK1* and *PARK2*^36^ (**Fig. 4I**).

**Figure 4.**
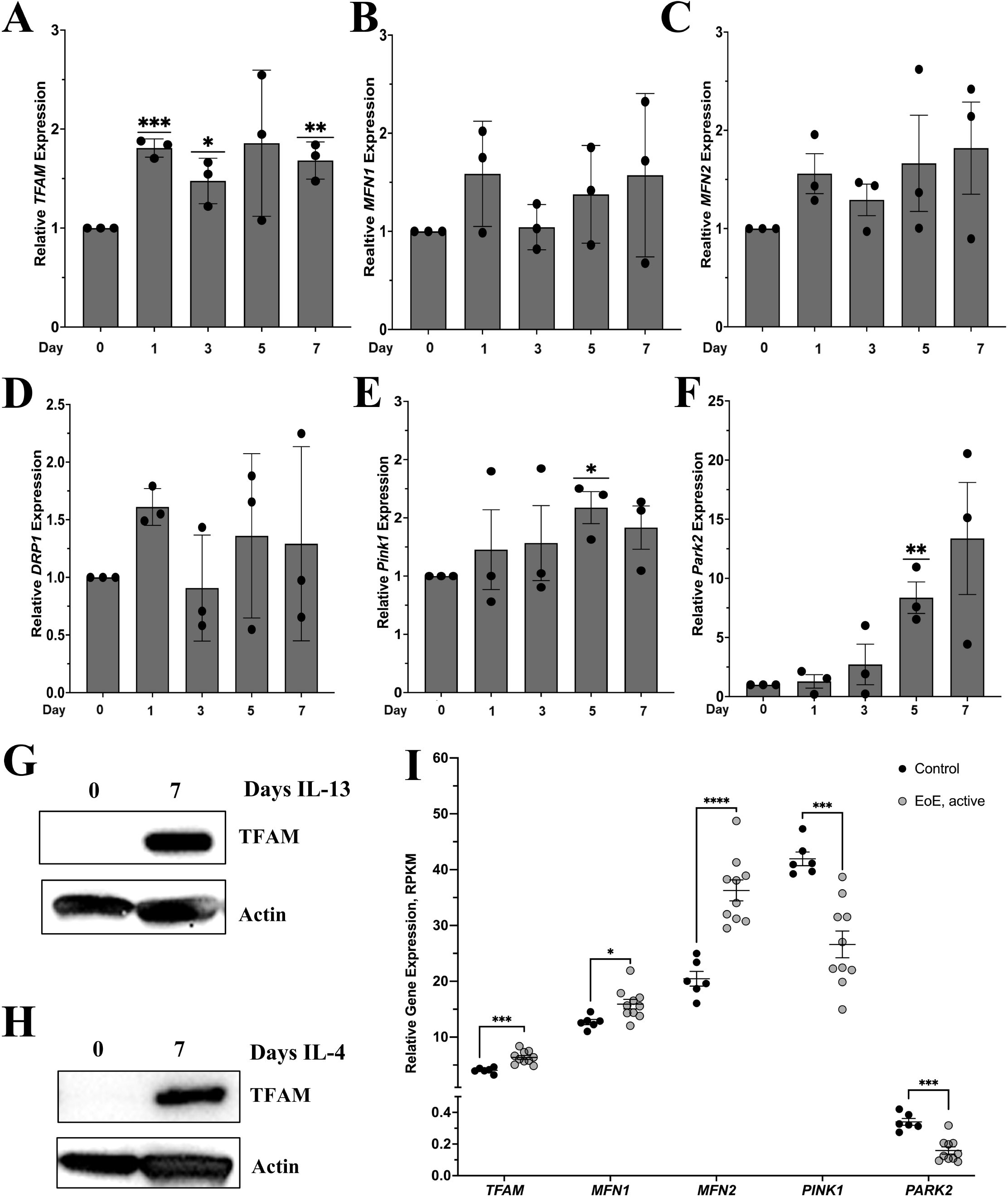
Effects of IL-13 and IL-4 on mediators of mitochondrial dynamics in esophageal keratinocytes. (**A-F**) EPC2-hTERT cells were treated with IL-13 (10 ng/mL) for the noted time points. Relative expression of (**A**) *TFAM,* (**B**) *MFN1,* (**C**) *MFN2,* (**D**) *DPR1,* (**E**) *PINK1,* and (**F**) *PARK2* was determined by qPCR. (**G, H**) Western blot showing for TFAM in EPC2-hTERT cells after 7 days of treatment with 10 ng/mL IL-13 (**G**) or IL-4 (**H**) with Actin as a loading control. (**I**) RNA expression of *TFAM, MFN1, MFN2, PINK1,* and *PARK2* expression in esophageal epithelium of active EoE patients and normal controls using data from Sherrill et al^36^. qPCR data was normalized to β-actin. Data in bar graphs presented as mean ± SEM, *p<0.05; **p<0.01; ***p<0.001; n=3.

### Pharmacological JAK/STAT inhibition limits IL-13-mediated increase in mtDNA and TFAM expression

As IL-13 and IL-4 activate JAK/STAT to promote eosinophilic infiltration and epithelial remodeling in EoE^16^, we next aimed to determine if alterations in mitochondrial biology occurring in esophageal epithelia cells responding to IL-13 or IL-4 are dependent upon JAK/STAT signaling. JAK inhibitor ruxolitinib effectively blocked STAT6 phosphorylation at Tyrosine residue 641 (**Fig. 5A**), the site required for STAT6 translocation to the nucleus^39, 40^. Ruxolitinib limited IL-13-mediated increases in both mtDNA and TFAM expression in EPC2-hTERT cells (**Fig. 5A, B**). IL-4-mediated induction of TFAM, however, was not affected by ruxolitinib (**Fig. 5A**).

**Figure 5.**
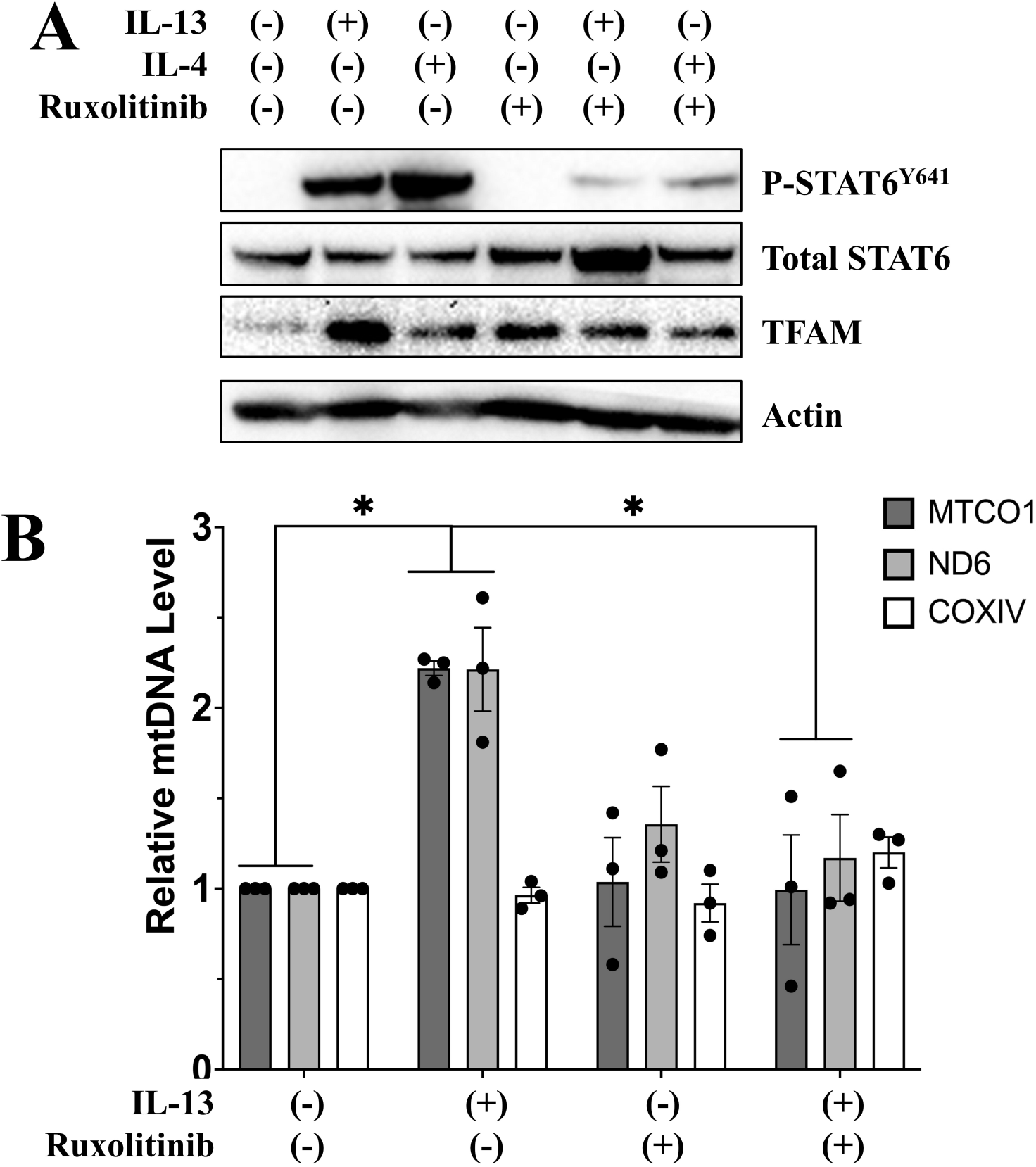
Effects of JAK inhibitor ruxolitinib on IL-13- and IL-4-mediated mitochondrial biogenesis in esophageal keratinocytes. EPC2-hTERT cells were treated with 10 ng/mL IL-13 or IL-4 for 7 days in the presence or absence of JAK inhibitor Ruxolitinib (100 ng/mL ). (**A**) Immunoblotting was used to assess expression of indicated proteins, including STAT6 phosphorylated tyrosine (Y) residue 641. Actin serves as a loading control. (**B**) Levels of mtDNA-encoded genes *MTCO1* and *ND6*, and nuclear-encoded gene *COXIV* were evaluated by qPCR. Data in bar graphs presented as mean ± SEM. *p<0.05; n=3.

### TFAM depletion limits IL-13-mediated esophageal epithelial remodeling in 3D esophageal organoids

IL-13 is a primary driver of epithelial remodeling in EoE^22, 41, 42^. As such, we next aimed to explore the functional role of IL-13-mediated increase in mitochondria upon esophageal tissue architecture. To do so, we generated 3D esophageal organoids from *Tfam^loxp/loxp^* mice (**Fig. 6A**). In the absence of Cre-mediated recombination, IL-13 induced a phenotype consistent with BCH, including impaired SCD, along with robust expression of TFAM (**Fig. 6B**). Upon *ex vivo* Cre-mediated recombination, IL-13-treated organoids from *Tfam^loxp/loxp^* mice displayed effective depletion of TFAM coupled with some evidence of SCD, including decreased basaloid cell content and the presence of centrally-localized keratinized core (**Fig. 6B**).

**Figure 6.**
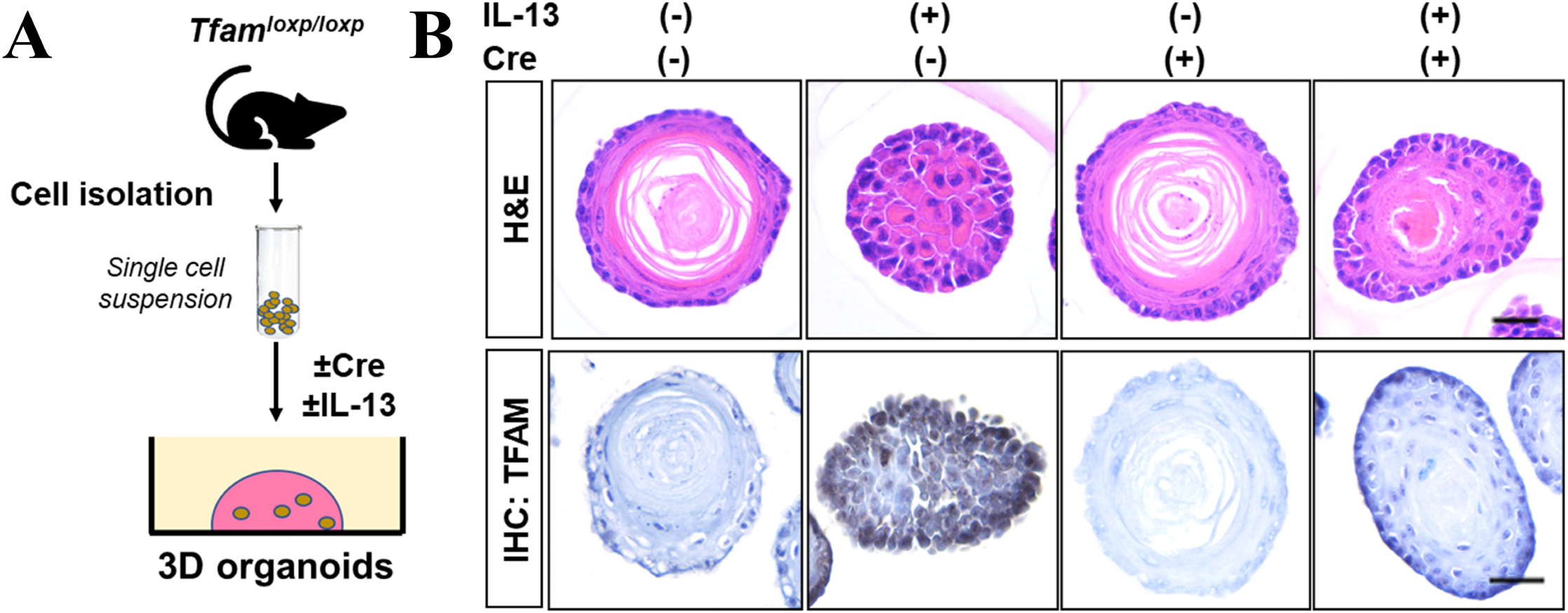
Effects of genetic depletion of TFAM on IL-13-mediated impairment of squamous cell differentiation in 3D organoids. (**A**) Schematic outlining approach for testing effects of Cre-mediated depletion of TFAM in organoids generated from esophageal epithelium of *Tfam^loxp/loxp^*mice. Adenoviral Cre and IL-13 (10 ng/mL) was added to organoids at the time of plating. After 7 days of growth, organoids were fixed and paraffin embedded. (**B**) Morphology of esophageal organoids was assessed via Hematoxylin & Eosin (H&E) staining and TFAM expression was assessed by IHC.

### IL-13 inhibits mitochondrial metabolism without impacting membrane integrity in esophageal keratinocytes

Recently, the bioenergetics in differentiated epithelium were shown to differ from that of undifferentiated epithelium in the esophagus^30^. As IL-13 limits SCD, we continued to examine the influence of IL-13 upon the metabolic profile of esophageal keratinocytes. Seahorse respirometry revealed a decrease in oxygen consumption rate of IL-13-treated EPC2-hTERT cells with suppression of basal, ATP-linked, and maximal respiration (**Fig. 7A-D**). Diminished energy production was further supported as EPC2-hTERT cells and primary esophageal epithelial cultured derived from either a normal non EoE control or patients with active EoE displayed decreases basal ATP levels upon IL-13 stimulation (**Fig. 7E-G**). To determine if decreased mitochondrial respiration is associated with mitochondrial membrane depolarization, we assessed MitoTracker Red (membrane potential sensitive dye) and MitoTracker Red Green (membrane potential insensitive dye) staining in IL-13 treated EPC2-hTERT cells. In these experiments, we found no evidence of alerted levels of mitochondrial depolarization in IL-13-treated cells (**Fig. 7H, I**).

**Figure 7.**
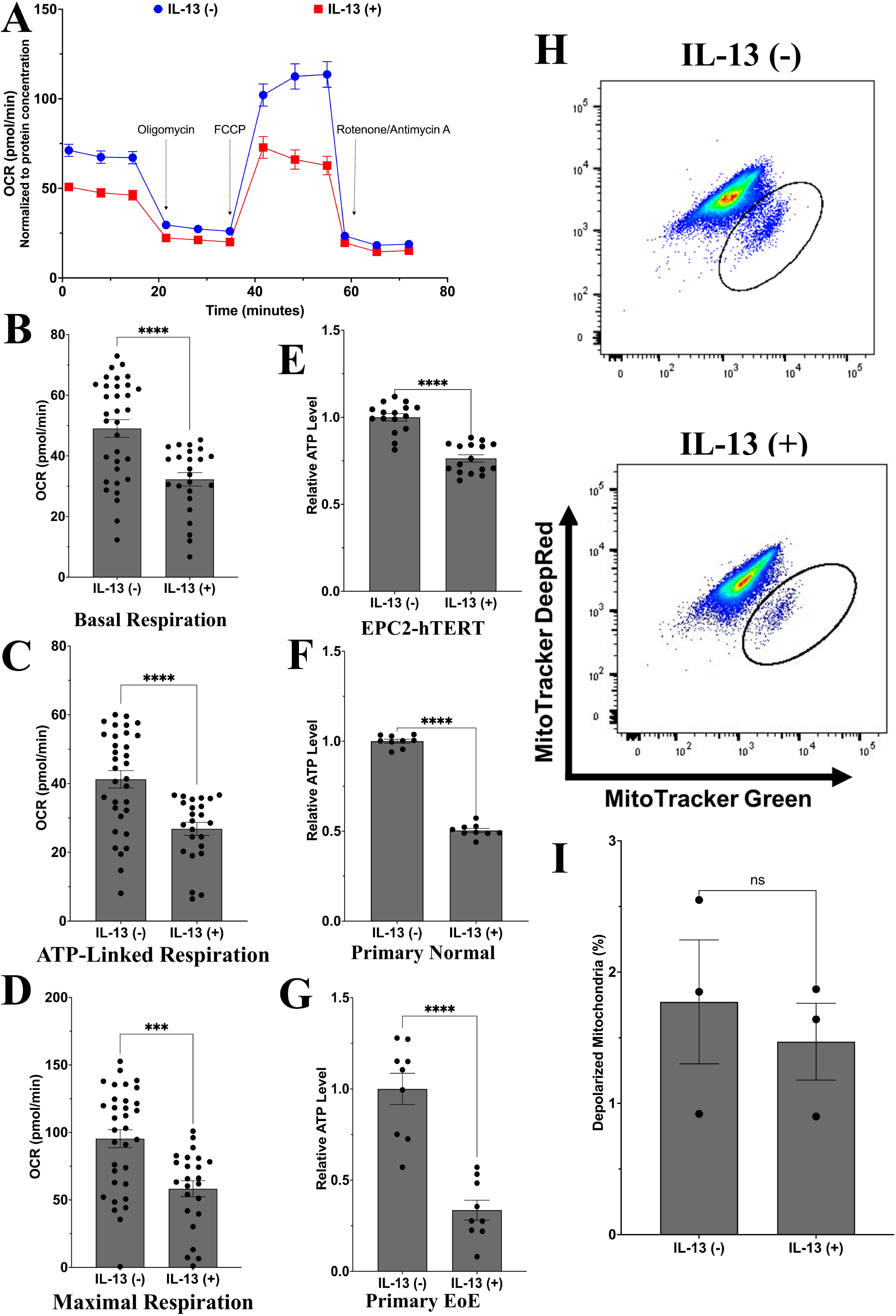
Effects of IL-13 on mitochondrial respiration, ATP level, and mitochondrial depolarization in esophageal keratinocytes. Esophageal epithelial cells were treated with IL-13 (10 ng/mL) for 7 days. (**A-D**) In EPC2-hTERT cells, mitochondrial bioenergetics was assessed by Seahorse respirometry. Specific parameters determined were (**A**) oxygen consumption rate; (**B**) basal respiration; (**C**) ATP-linked respiration; and (**D**) maximal respiration. (**E-G**) Relative ATP measurement by ATP determination assay in (**E**) EPC2-hTERT cells; (**F**) normal primary esophageal keratinocytes; (**G**) and (**G**) primary esophageal keratinocytes derived from patient with active EoE. (**H, I**) MitoTracker Red/Green flow cytometry was performed in EPC2-hTERT cells. (**H**) Representative flow cytometry dot plots. (**I**) Bar diagram showing percentage of cells with depolarized mitochondria. Data in bar graphs presented as mean ± SEM. *p<0.05; **p<0.01; ***p<0.001; ****p<0.0001; n=3.

### IL-13 suppresses mitochondrial reactive oxygen species, and this effect is blocked by TFAM depletion

Finally, as ROS have been shown to induce SCD in esophageal epithelial cells^21^ and mitochondria are a primary source of ROS, we investigated the relationship between IL-13 and oxidative stress in EPC2-hTERT cells. We found that IL-13 limited levels of mitochondrial superoxide in EPC2-hTERT cells (**Fig. 8A, B**). siRNA-mediated TFAM depletion also limited mitochondrial superoxide levels in the absence of IL-13 stimulation (**Fig. 8C, D**); however, when EPC2-hTERT cells were depleted of TFAM, IL-13 treatment failed to limit mitochondrial superoxide levels (**Fig. 8C, D**).

**Figure 8.**
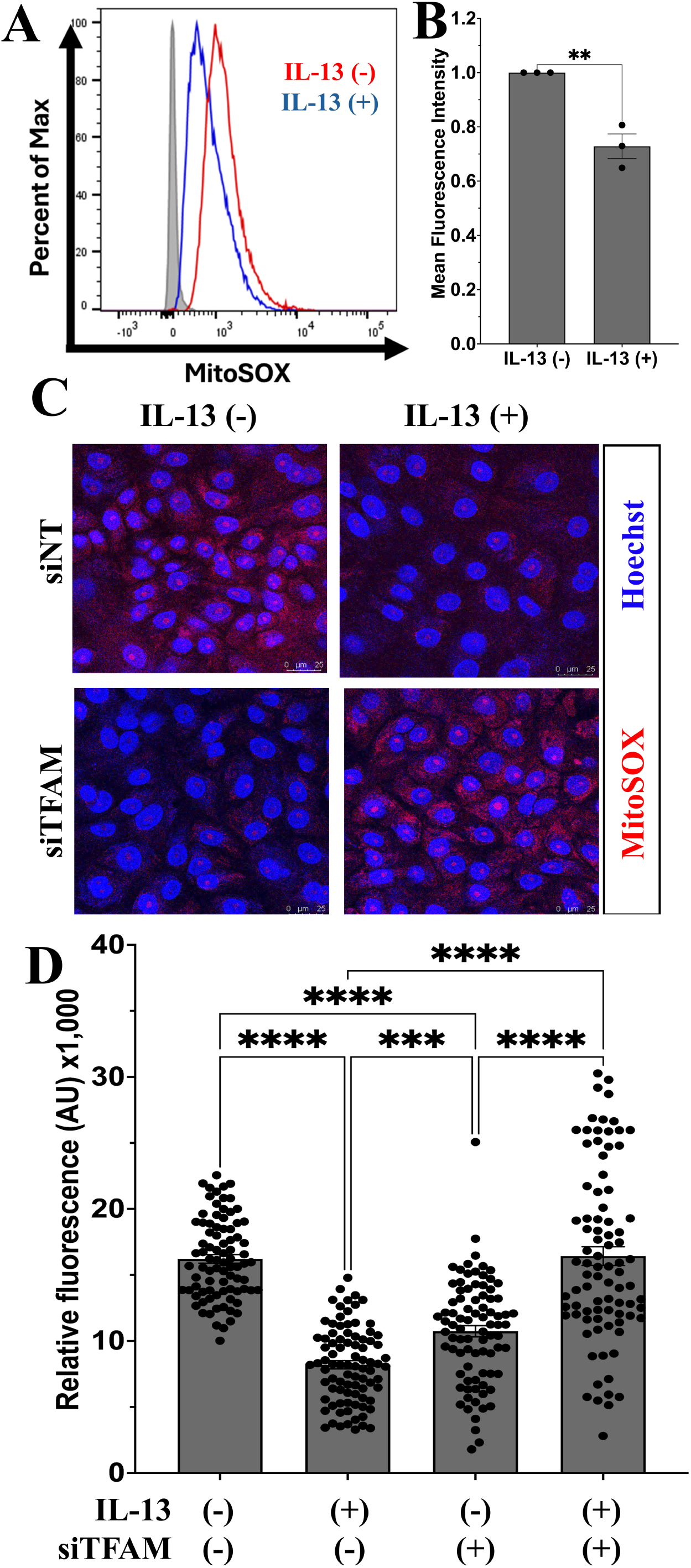
Effects of TFAM depletion on IL-13-mediated suppression of mitochondrial reactive oxygen species. (**A, B**) EPC2-hTERT cells were treated with 10 ng/mL IL-13 for 7 days then stained with MitoSOX fluorescence and assessed by flow cytometry. (A)of IL-13-treated EPC2-hTERT cells with mean fluorescent intensity bar graph. (**A**) Representative flow histograms and (**B**) bar diagram showing average mean fluorescence intensity are shown. (**C, D**) EPC2-hTERT cells were transfected with siRNA targeting TFAM (siTFAM) or non-targeting (siNT) control oligonucleotides for 72 hours. Cells were then cultured in the presence or absence of IL-13 (10 ng/μl) for 7 days and stained with MitoSOX Red and Hoechst 33342 to assess superoxide and nuclei, respectively, by confocal imaging. (**C**) Representative confocal images and (**D**) bar diagram quantifying relative fluorescence are shown. Data in bar graphs presented as mean ± SEM. **p<0.01; ***p<0.001; ****p<0.0001; n=3.

## Discussion

While mitochondria have been implicated in various immune-mediated inflammatory pathologies, including asthma and eczema^43, 44^, our understanding of the role of this organelle in EoE pathogenesis is limited. We initiated our study by investigating levels of the mitochondrial protein MTCO1 in esophageal biopsy specimens from EoE patients as well as non-EoE controls. We observed a significant increase in the expression of MTCO1, subunit of mitochondrial complex IV, in active EoE patients as compared to either subjects with inactive EoE or controls. We further identified an increase in MTCO1 expression and mtDNA levels in esophageal epithelium of mice with MC903/OVA-induced EoE-like inflammation. In agreement with our findings, a recent report found increased expression of a marker of mitochondrial complex V (ATP synthase) in esophageal epithelium of patients with active EoE^30^. Additionally, although Sherril and colleagues reported decreased levels of DHTKD1 RNA and protein in familial EoE cases, an opposite trend was seen in non-familial EoE cases^27^. Heterogeneity in MTCO1 IHC score was present within each group of human subjects and mice in our datasets. As our *in vitro* studies suggest that IL-13 and IL-4 promote increased mitochondrial mass in esophageal keratinocytes, future investigations will determine if high levels of MTCO1 may be reflective of a subset of EoE patients who are more likely to respond to dupilumab, serving as a predictive biomarker of response. Such investigations will be executed in a prospective and longitudinal fashion, in contrast to the current study in which MTCO1 expression was evaluated retrospectively in a cross-sectional cohort. Notably, metabolic alterations, including activation of the mitochondrial tricarboxylic acid cycle, have been documented in atopic dermatitis patients classified as good responders to dupliumab as compared to poor responders^45^. As mitochondrial dysfunction has been linked to fibrosis in a variety of tissues^46^, evaluation of the relationship between mitochondrial mass in esophageal epithelium and EoE phenotypes in mice and endotypes in human subjects^47^ is warranted.

To investigate the influence of EoE inflammatory mediators on mitochondrial mass, we treated esophageal epithelial cells with a panel of EoE-relevant cytokines. Cytokine treatments were performed over 7 days as this was the earliest time point at which reproducible changes in mtDNA content were observed in esophageal epithelial cells *in vitro* (KAW, unpublished data). We found that Th2 cytokines IL-13 and IL-4 significantly increased mitochondrial mass while TNFα significantly decreased mitochondrial mass. As we observed evidence of increased mitochondria in esophageal epithelium of humans and mice with EoE inflammation, the current study investigated the influence of IL-13 and IL-4 on mitochondria in esophageal epithelial cells. However, as the EoE inflammatory milieu is complex, future studies utilizing pharmacologic and/or genetic approaches in murine EoE models will be informative with regard to the inflammatory signals that increase mitochondrial mass in esophageal epithelium *in vivo*. As TNFα is a driver of EoE-associated fibrosis^48–50^, it is possible that the balance between IL-13/IL-4 and TNFα and the resulting changes in mitochondrial mass may influence whether EoE presents as inflammatory or fibrotic, which could be tested in murine EoE models. As our own work has demonstrated that aged mice more readily exhibit fibrosis as compared to their young counterparts^51^, it will be of interest to examine mitochondrial levels and expression of EoE-relevant cytokines in young and aged mice with EoE-like inflammation. Here, we report that IL-13 and IL-4 induce the expression of TFAM, a critical mediator of mitochondrial biogenesis. As TNFα has been shown to suppress mitochondrial biogenesis concomitant with downregulation of TFAM expression in fat and muscle of obese rodents^52^, it is possible that TNFα may inhibit mitochondrial biogenesis in esophageal keratinocytes. Alternatively, TNFα may influence other aspects of mitochondrial biology to decrease mitochondrial mass. We have previously demonstrated that both IL-13 and TNFα promote activation of autophagic flux in esophageal keratinocytes^18^. As mitophagy is a mechanism to clear mitochondria from cells, it is possible that the balance between mitophagy and mitochondrial biogenesis is skewed toward biogenesis in IL-13-treated esophageal keratinocytes while the opposite may be true in cells responding to TNFα.

Our data demonstrate that JAK/STAT signaling contributes to the induction of TFAM in esophageal epithelial cells responding to IL-13. To the best of our knowledge, STAT-mediated transcriptional regulation of TFAM has not been reported. STAT3 has, however, been shown to interact with TFAM at mtDNA, contributing to regulation of mitochondrially-encoded gene expression in murine epidermal keratinocytes^53^. In this context, mitogen signaling phosphorylates STAT3 at Serine 727, permitting STAT3 localization to mitochondria where it interacts with mtDNA and TFAM to limit expression of mitochondrially-encoded genes^53^. While STAT3 depletion augmented mitochondrial respiration in murine epidermal keratinocytes under basal conditions, no effect on mtDNA level was identified^53^. Consistent with this report, we found that pharmacological JAK inhibition had no effect on mtDNA at baseline. We did, however, see that ruxolitinib increased expression of TFAM at the protein level in esophageal keratinocytes in the absence of cytokine stimulation. IL-6-mediated activation of STAT3 has been linked to TFAM downregulation in colonic epithelial cells via microRNA-23b^54^. Thus, there may be multiple mechanisms through which STATs regulate TFAM in a cell-type and/or context-dependent manner. Although recent evidence supports a role for STAT3 in EoE pathogenesis^42^, STAT6 is well-established to contribute to EoE pathobiology, including by promoting transcriptional upregulation of eotaxin-3^15, 16, 55^, follistatin^21^, calpain-14^14, 23, 56^, and synaptopodin^24^ in esophageal epithelial cells responding to IL-13 or IL-4. Although STAT6 has been shown to translocate to mitochondria and regulate mitochondrial function and fusion in non-epithelial cells^57–60^, what role, if any, STAT6 plays in regulation of mitochondrial biology in epithelial cells remains to be determined. STAT1 has also been implicated in EoE^61^ and shown to localize to mitochondria and antagonize mitophagy in cardiomyocytes^62^. As we continue to study the regulation of mitochondria by IL-13 and IL-4, we will explore how these Th2 cytokines influence expression and localization (i.e. nucleus vs. mitochondria) of individual STATs as well as the effects of modulating STAT expression, localization, and interactions on mitochondrial biology.

In addition to identifying how the EoE inflammatory milieu influences mitochondrial mass in esophageal epithelium, the current study aimed to investigate the functional role of mitochondria in EoE pathogenesis. Using 3D organoids, we found that genetic depletion of TFAM partially restored IL-13-mediated impairment of differentiation. As excessive ROS has been shown to induce esophageal epithelial differentiation *in vivo*^21, 25^ and mitochondria are a primary source of cellular ROS, we also evaluated the relationship between IL-13, mitochondrial superoxide, and TFAM. In IL-13-treated esophageal epithelial cells, mitochondrial superoxide levels were suppressed. This is consistent with studies showing that although IL-13 initially induces ROS in esophageal epithelium this oxidative stress is alleviated as mechanisms to counteract it are activated, including autophagy and expression of antioxidants^18, 21^. We also find that mitochondrial respiration, which produced ROS as a byproduct, is diminished in IL-13-treated esophageal keratinocytes. It is possible that increased mitochondrial mass occurs as a compensatory mechanism to support energy demands in IL-13-treated cells. Time course experiments will help to clarify the relationship between alterations in mitochondrial mass, respiratory activity, and oxidative stress. The relationship between TFAM and oxidative stress remains ambiguous with some studies showing increased ROS with TFAM knockdown while others demonstrate decreased ROS^63–68^. In EPC2-hTERT cells genetic depletion of TFAM limited mitochondrial superoxide under basal conditions. In contrast, superoxide levels in IL-13-treated esophageal cells were restored upon depletion of TFAM, which also partially restores SCD in IL-13-treated esophageal organoids. The mechanisms underlying the differential effects of TFAM depletion on mitochondrial ROS in esophageal keratinocytes in the presence and absence of IL-13 remain to be determined.

One final consideration is examination of how IL-13-mediated increase in mitochondrial mass in esophageal keratinocytes may relate to proliferation, which is coupled with SCD in esophageal epithelium wherein proliferative basal/suprabasal cells exit the cell cycle as they differentiate. Diminished reliance on mitochondrial metabolism is a feature of many dividing mammalian cells that preferentially use glycolysis for energy production as it increases availability of carbon and other resources that are needed to increase biomass^69^. Although this concept provides rationale to support diminished mitochondrial activity in IL-13-treated esophageal keratinocytes, it also operates under the assumption that IL-13 drives proliferation in this context. The reality may be more complex. In EPC2-hTERT cells, heterogeneity was recently identified with regard to proliferation as clonal tracing revealed that only ∼10% of clones exhibit exponential growth in non-genetic heritable fashion^70^. Thus, effects on mitochondria and cell metabolism mediated by EoE-relevant cytokines may represent behaviors attributable to subsets of cells, which could be explored using single-cell based approaches. Evaluation of cellular heterogeneity in esophageal epithelium and in response to inflammatory stress may also help to reconcile our findings with a recent study indicating that enhanced mitochondrial activity is associated with impaired differentiation in esophageal epithelium^30^. In the study by Ryan et al, dysregulated hypoxia-inducible factor (HIF)-1α signaling enhances mitochondrial respiration and diminishes glycolysis which is associated with impaired calcium-induced differentiation *in vitro*^30^. In macrophages, HIF-1α-mediated induction of glycolysis has been shown to promote classical pro-inflammatory activation, while IL-4-mediated induction of STAT6 and mitochondrial biogenesis supports alternative anti-inflammatory activation. These findings supporting the premise that integration of cell signaling and metabolism may influence specific cell fate decisions in a cell population. With regard to the role of proliferation in EoE-associated epithelial remodeling, biopsies from patients with active EoE exhibit an increase in both SOX2-postive and KI67-positive cells; however, no significant difference in proliferative (Ki67-positive) basal (SOX2-poitive) cells was found in these subjects^17^. In both EoE patients and mice with MC903/OVA-mediated EoE-like inflammation, reported evidence of suprabasal cell expansion has been reported^71, 72^, indicating that BCH in EoE may be linked to factors beyond an increase in basal cell proliferation. Supabasal cell expansion was also identified in human subjects with EoE^72^. In mice, EoE-associated suprabasal cells exhibited evidence of senescence^71^, a pathway that is linked to mitochondrial ROS production. In salivary epithelial cells, STAT6 signaling has been linked to mitochondrial dysfunction and ROS production that drives senescence^59^.

In conclusion, we utilized human and murine esophageal tissue with EoE inflammation as well as *in vitro* and *ex vivo* models to delineate the role of mitochondria in EoE pathogenesis. Specifically, IL-13 promotes increased intracellular mitochondria within the esophageal epithelium which in turn contributes to impaired squamous differentiation. Moreover, diminished ROS in response to IL-13, a known regulator of differentiation, is supported by increased mitochondrial biogenesis. We propose a model wherein IL-13-mediated STAT signaling promotes expression of TFAM and accumulation of mitochondria with diminished respiratory activity, providing a limited supply of ROS to induce SCD. Thus, these findings provide novel insights into the underlying mechanisms through which EoE inflammatory mediators influence mitochondrial biology as well as the functional role of mitochondria in EoE pathogenesis.

## Abbreviations

ATP: adenosine triphosphate
DHTKD1: dehydrogenase and transketolase domain containing 1
EGD: esophagogastroduodenoscopy
EGID: eosinophilic gastrointestinal diseases
EoE: eosinophilic esophagitis
eos/HPF: eosinophils per high power field
ETC: electron transport chain
FCCP: carbonilcyanide p-triflouromethoxyphenylhydrazone
H&E: hematoxylin & eosin
HIF: hypoxia-inducible factor IL: Interleukin
IL-4R: IL-4 receptor
KSFM: keratinocyte serum free media
MTCO1: mitochondrially encoded cytochrome c oxidase I
mtDNA: mitochondrial DNA
OCR: oxygen consumption rate
OGDHL: Oxoglutarate dehydrogenase-like
OVA: ovalbumin
OXPHOS: oxidative phosphorylation
ROS: reactive oxygen species
SCD: squamous cell differentiation
siNT: siRNA non-targeting
STAT: signal transducer and activator of transcription
TFAM: transcription factor a, mitochondria
Th: T helper
TNF: Tumor necrosis factor

## Acknowledgments

We thank the following former members of the Whelan lab for their support Anbin Mu, Timothy Hall, Anne-Laure Monéger, Julie Gang, and M. Faujul Kabir. We acknowledge the staff of following core facilities for technical assistance: Temple University Lewis Katz School of Medicine Flow Cytometry Core, (Director, Amir Yarmahoodi); Fox Chase Cancer Center Histopathology Core; Children’s Hospital of Philadelphia Gastrointestinal Epithelium Modeling Program Core (RRID: SCR_026402) and the University of Pennsylvania Center for Molecular Studies in Digestive and Liver Diseases (NIHP30DK050306).

