## Supplementary figures and images for "Interleukin-13-mediated alterations in esophageal epithelial mitochondria contribute to tissue remodeling in eosinophilic esophagitis"

### Figure E1

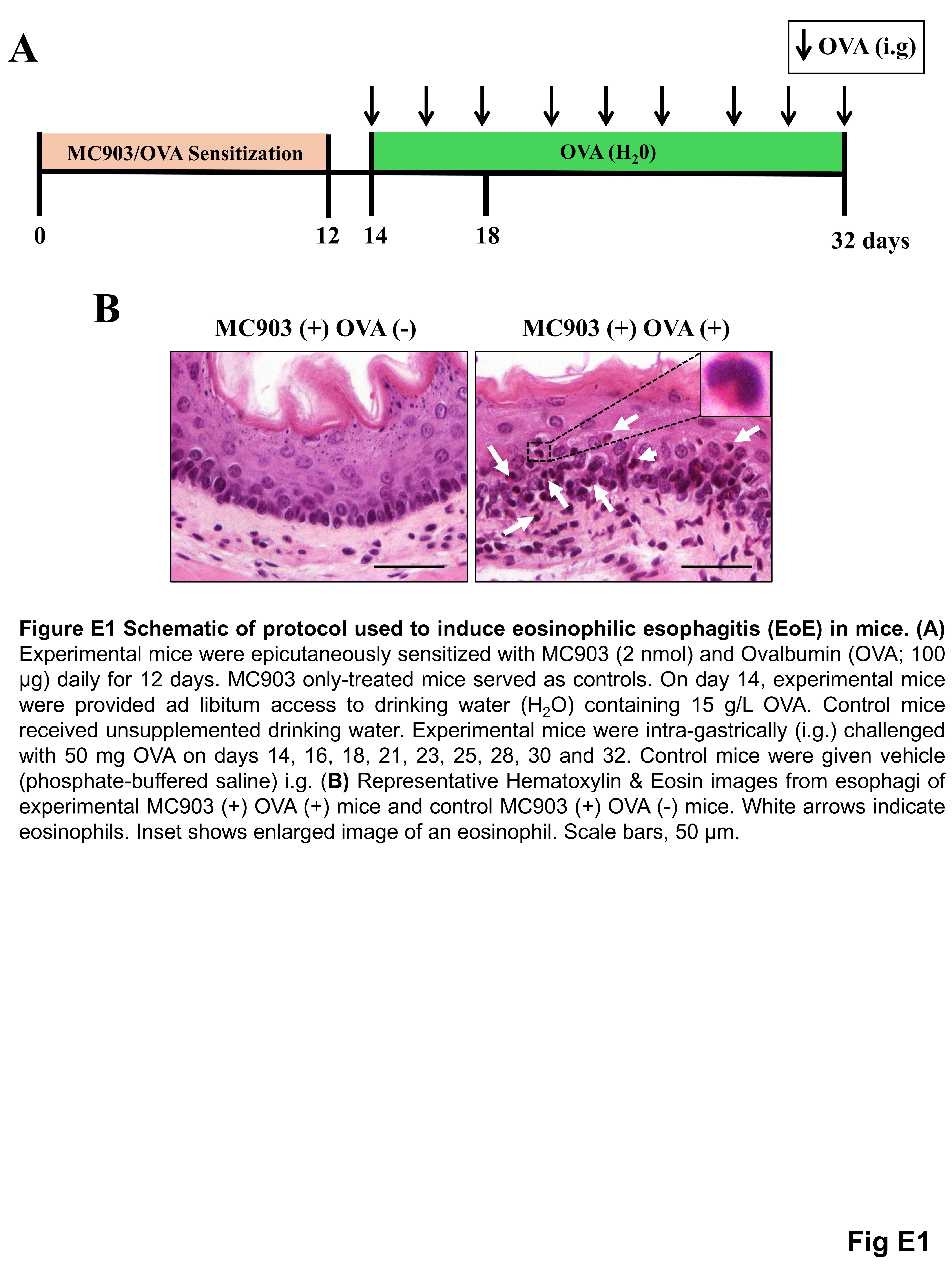

### Figure E2

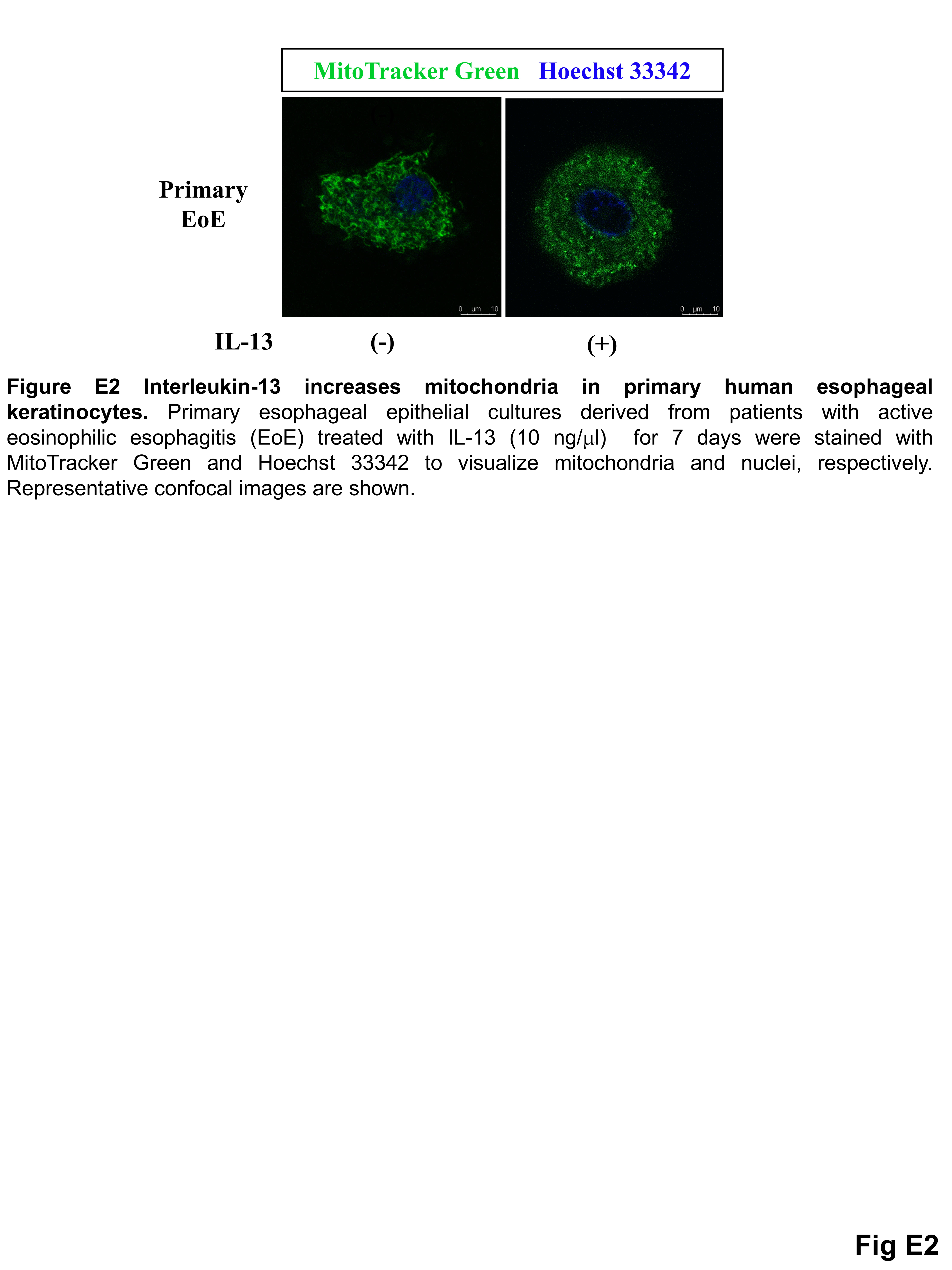

### Figure E3

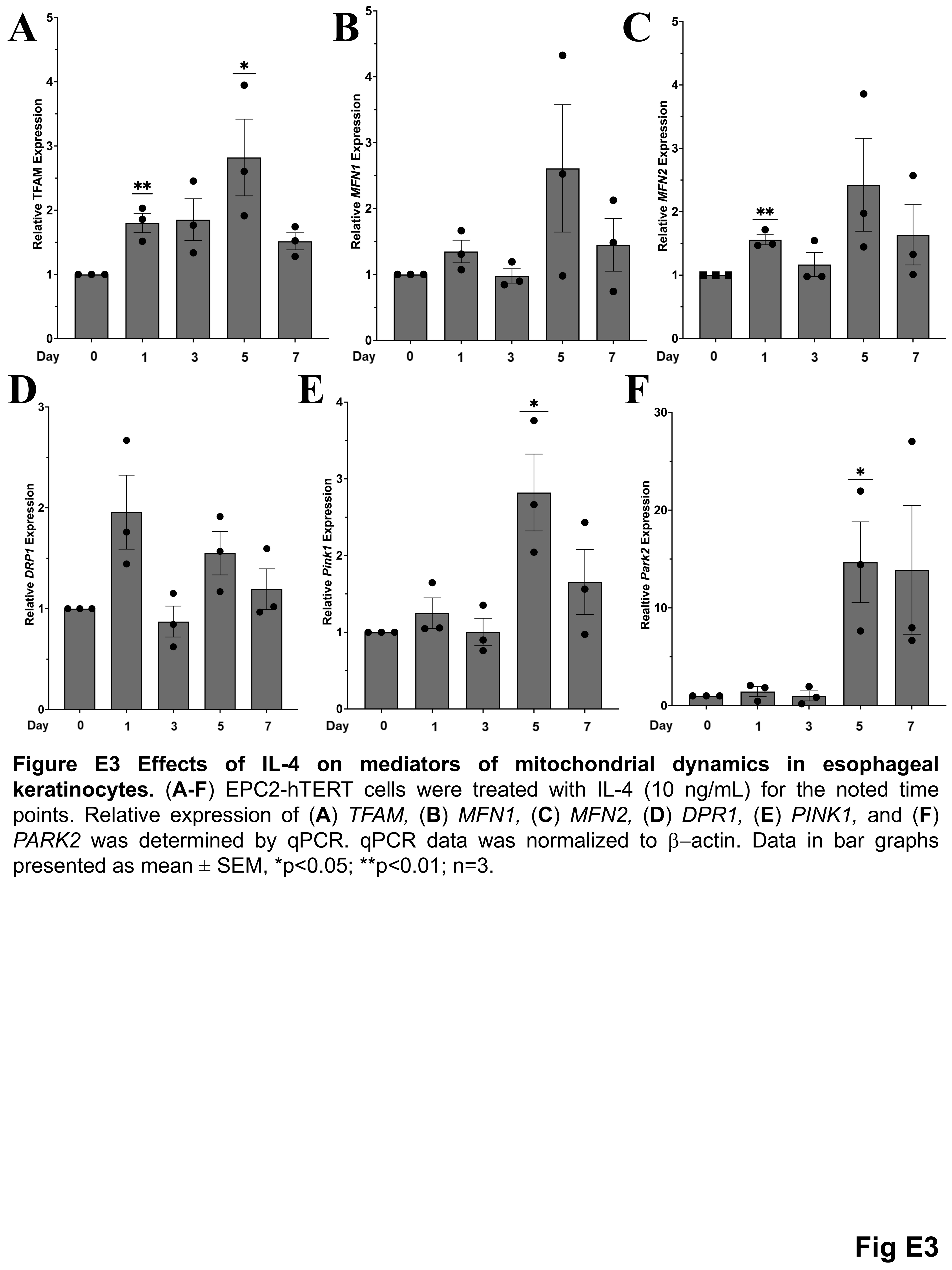

### Table E1

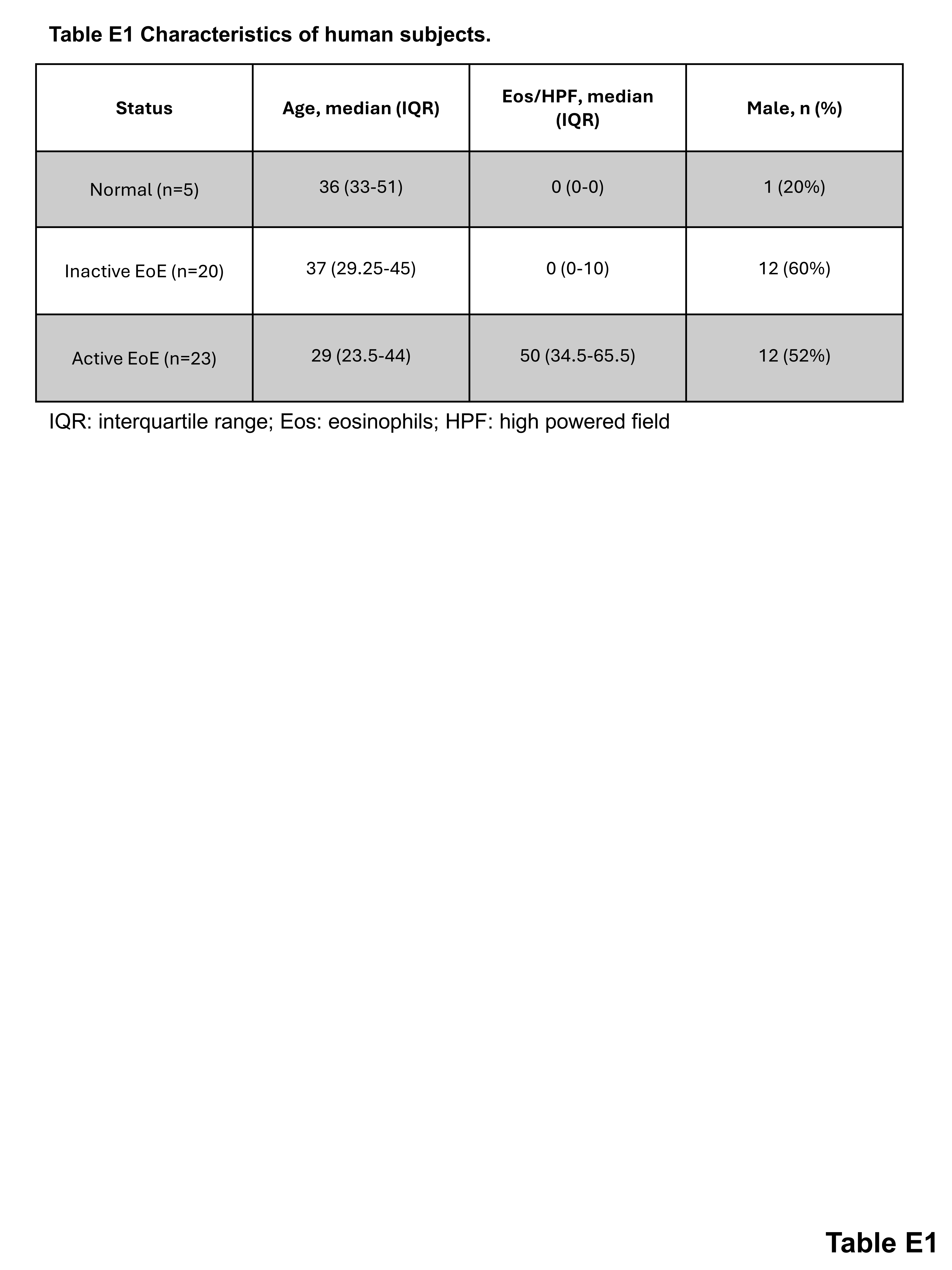

### Table E2

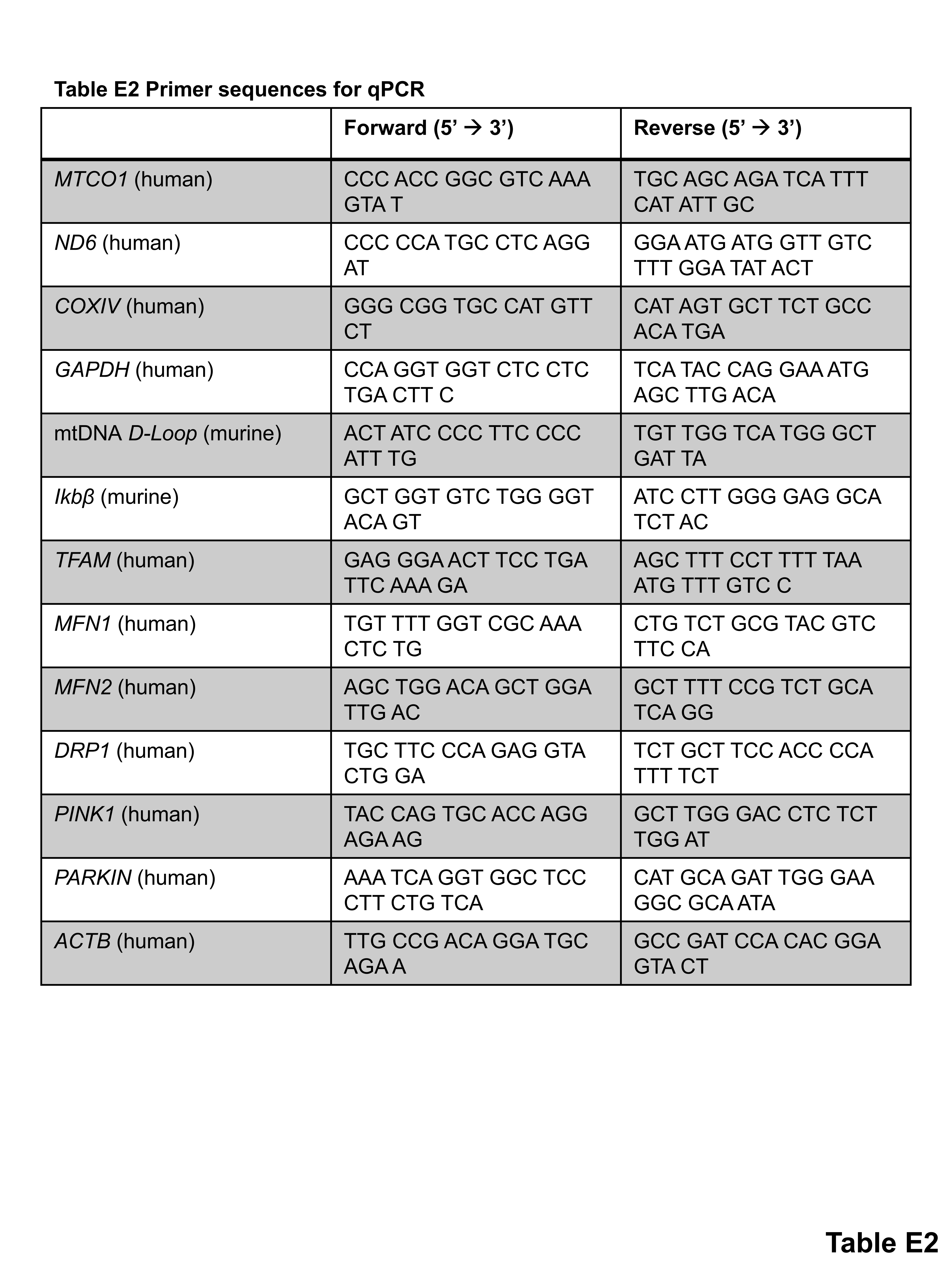
